# A combinatorial extracellular code tunes the intracellular signaling network activity to distinct cellular responses

**DOI:** 10.1101/346957

**Authors:** Dmitry Kuchenov, Frederik Ziebell, Florian Salopiata, Mevlut Citir, Ursula Klingmueller, Wolfgang Huber, Carsten Schultz

## Abstract

Cells constantly survey a complex set of inputs that is processed by the intracellular signaling network, but little is known of how cells integrate input information from more than one cue. We employed a FRET biosensor-based imaging platform to study the effect of combinatorial growth factor levels on the signaling network in human cells. We found that pairwise stimuli caused distinct concentration- and ratio-dependent signaling states through signaling signatures such as antagonism, additivity and synergy. The unique signaling states correlated with differential gene expression and non-additive transcription patterns. We further elucidated how a signal-rich environment can fine-tune the signaling network and adjust physiological outcomes, by kinase and phosphatase activity profiling. We describe how complex extracellular conditions affect phospho-turnover and the basal phosphorylation status. Thus, we provide mechanistic insights into cellular processing of multiple cues and explain part of the complexity of cellular adaptation to changes in the extracellular environment.

## INTRODUCTION

An extracellular microenvironment is a highly signal-rich system that is partly characterized by the identity and concentration of signaling cues such as growth factors (GFs) and cytokines (Watson et al., 2018). Cells in an organism continuously monitor their dynamic microenvironment to make decisions on survival, growth, proliferation, differentiation, secretion or migration (Rezza et al., 2016; Sever and Brugge, 2015). Remarkably, such cellular function is achieved through a limited number of intracellular signaling components to process most of the extracellular cues (Ammeux et al., 2016). The components are organized in an intricate network of dynamic physical interactions and biochemical reactions (Wilkes et al., 2015). In particular, the dynamic properties of the molecular events seem to be employed by cells to accurately process information from the extracellular environment (Purvis and Lahav, 2013; Selimkhanov et al., 2014; Sonnen and Aulehla, 2014). Although aspects of network architecture and signaling dynamics have been the subject of intensive study (Bakal et al., 2008; Hill et al., 2016; Selimkhanov et al., 2014; Wagner et al., 2013; Wilson et al., 2017), at the systems level it remains unclear how cells integrate and process information from multiple external cues.

Receptor tyrosine kinases (RTKs) are well known to employ combinatorial signalling pathways to govern diverse fate decisions and biological processes such as proliferation, apoptosis, differentiation and metabolism. Aberrant RTK signaling is frequently involved in diseases including cancer (Verstraete and Savvides, 2012; Wilson et al., 2012). A ligand, such as a growth factor (GF) or cytokine, binds to an RTK at the plasma membrane inducing signaling through a set of well-known intertwined signalling pathways that are shared among different RTK signalling networks (Hill et al., 2016; Kolch et al., 2015; Lemmon and Schlessinger, 2010; Wagner et al., 2013). Yet, different RTKs have quite distinct functions *in vivo* (Chao, 1992; Vasudevan et al., 2015). Remarkably, combinations of GFs/cytokines may induce a unique gene expression profile as well as a physiological outcome (Beaujean et al., 2003; Cappuccio et al., 2015; Martin et al., 2009). We therefore hypothesized that studying the dynamic aspects of GF signaling in a combinatorial context in living cells may reveal new clues about the function of the underlying signaling network.

In previous studies, putative mechanisms of signal integration have been identified within a signaling network using different cells and conditions but so far no clear overarching model has been established. Various reports propose that signaling cross-talk or feedback loops between and within pathways result in signaling interactions (or cross-talk) such as additivity, synergy and antagonism. (Ammeux et al., 2016; Borisov et al., 2009; Natarajan et al., 2006). Our recent work has suggested that the relative level and concentration of GFs in a pairwise stimulation will program signaling signatures (Kuchenov et al., 2016). Such signatures might be unique and are not predictable from the individual components of a signaling network and/or the individual treatments with a single cue which makes signaling network analysis a challenge (Borisov et al., 2009; Chatterjee et al., 2010; Hsueh et al., 2009). However, a combinatorial stimulation with simultaneous quantitative monitoring of multiple signaling network events over time would help to better understand the integration of information from multiple cues and subsequently the function of signaling networks in live cells. Such extended data sets will reflect the complexity of the cellular signaling networks, but are labor-intensive to generate in a comprehensive fashion by conventional approaches. To meet this challenge, we previously developed the FRET-based multi-parameter imaging platform (FMIP) that allowed monitoring the activity of 40 signalling molecules at the single cell level with high temporal resolution in a single experiment (**Figure 1A**) (Kuchenov et al., 2016). The design of the platform permits continuous probing the signalling of multiple components in living cells to numerous conditions under identical experimental settings in a very time-efficient manner.

**Figure 1.**
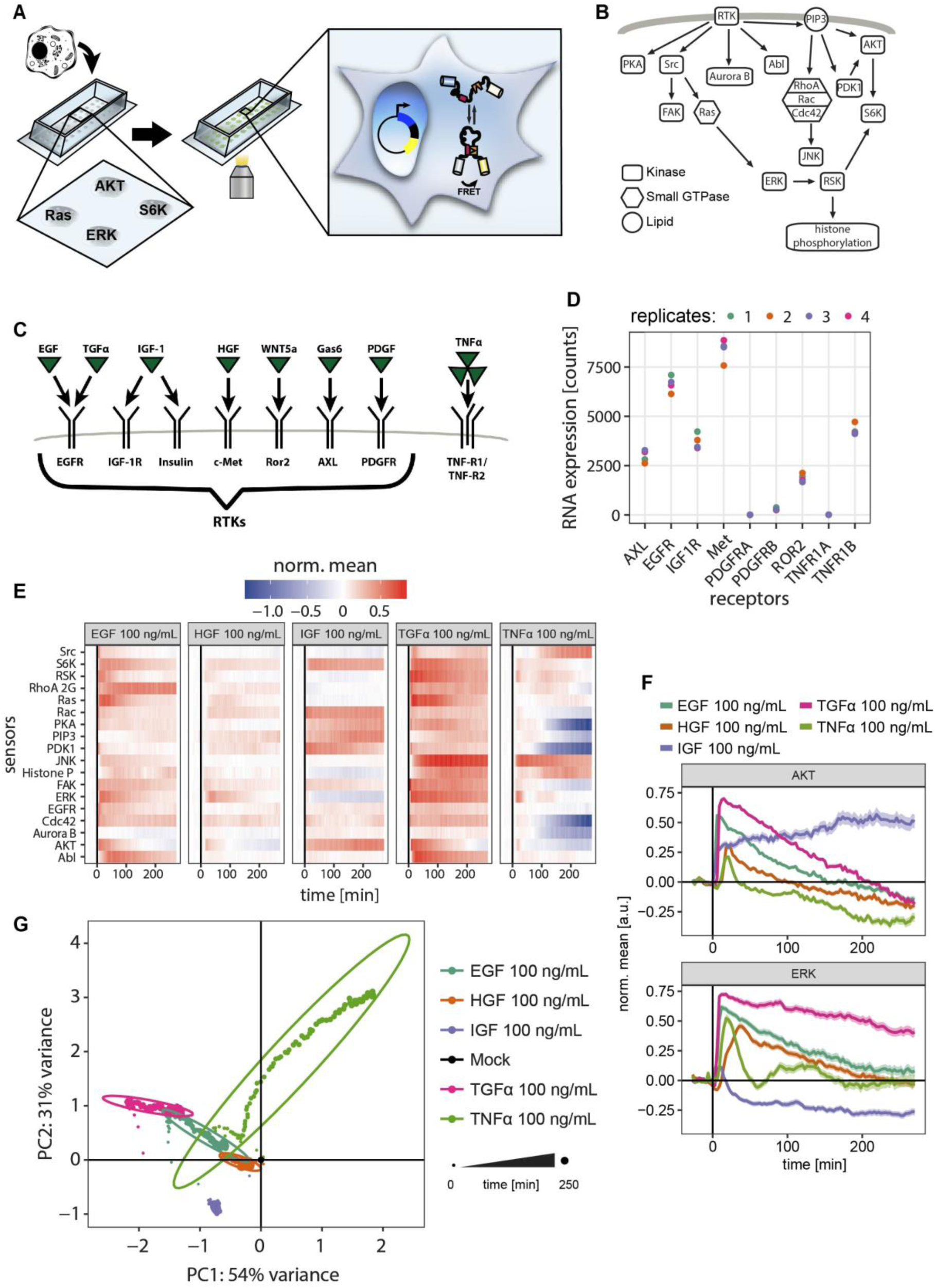
Dynamic signaling program is defined by cue identity. (**A**) Schematic depiction of FMIP assay. Plasmids encoding FRET biosensors were individually reverse transfected into human cells. FRET ratio was measured with a wide-field automated microscope. (**B**) Overview of the GF signaling network measured by the FMIP. (**C**) Schematic depiction of growth factor/cytokine signaling induction pursued in this work. (**D**) The expression level in HeLa cells of the investigated receptors. The abundance of mRNA was determined by RNA-seq after 16 hours of serum starvation. n = 4 independent experiments (**E**) Dynamic signaling response after exposure to 5 different ligands. Color scale indicates mean FRET ratio over 270 min, normalized by the absolute maximum value (relative activity). (**F**) Representative examples of distinct dynamics after GF or cytokine treatment for the data quantified in (D). Curves indicate mean ± SEM. (**G**) Principal component analysis (PCA) of the average response to growth factors and cytokines. Each dot represents a single time point while the ovals indicate 95% of time points of the same treatment.

In this work, we use the FMIP assay to quantify multiple signaling events upon stimulation with the pairwise input (EGF, IGF-1, HGF, TGFα, TNFα) of various doses and ratios. We show that such pairwise GF stimulation shapes the global signaling program through dynamic signaling signatures classified as additivity, synergy and antagonism. In turn, each signaling network states correlates with distinct gene expression profiles and transcription interactions. We further demonstrate that in the presence of serum, i.e. under quasi-physiological conditions, the cellular signaling network is pre-activated and tuned to achieve a unique signaling program in response to low concentrations of EGF. We report that the steady-state phospho-turnover rate and phosphorylation status of downstream RTK effectors differs qualitatively and quantitatively depending on the presence of multiple signaling cues. Our findings reveal that a complex environment defined by the identity, concentration and combinatorial ratio of the extracellular cues plays a crucial role in history-guided cellular adaptation to a changing environment.

## RESULTS

### Similarity and distinction in the TGFα, EGF, IGF-1, and HGF signalling programs

We employed a FRET biosensor-based multi-parameter imaging platform (FMIP) for monitoring 40 signalling events in near real-time at the single cell level with high temporal resolution in a single experiment (**Figure 1A**) (Kuchenov et al., 2016). This system relies on transient expression of FRET-biosensors that visualize enzyme activities in live cells rather than the protein posttranslational modification status (Kuchenov et al., 2016; Newman et al., 2011). Therefore, this platform quantitatively measures the functional status of a signalling network (Kuchenov et al., 2016).

To investigate integration mechanisms of multiple extracellular signals, we first set out to assess the signalling response induced by various growth factors. We stimulated HeLa cells with 100 ng/mL of platelet-derived growth factor BB (PDGF-BB), epidermal growth factor (EGF), tumor growth factor alpha (TGFα), insulin-like growth factor (IGF-1), hepatocyte growth factor (HGF), growth arrest-specific protein 6 (GAS6), wingless and int-related protein 5a (WNT5a) or tumor necrosis factor alpha (TNFα) and monitored responses of 40 FRET biosensors by the FMIP (**Figure 1A**). TNFα, which binds to the receptor of a TNFR protein family, was included to reveal general principles of signal integration. Each of these signaling molecules binds to a well-known receptor inducing signaling that has been intensively characterized before (**Figure 1B-C**) (Beyer and MacBeath, 2012; Wagner et al., 2013). Of the 40 FRET sensors, we identified 18 that reported consistent FRET ratio changes across biological replicates (n ≥ 2, 262 cells in average) under at least one condition. Collectively, these biosensors visualized a broad portion of the signalling network activity including Ras/ERK, PI3K/AKT, JNK, and Src/FAK pathways (**Figure 1C**). In these cells, PDGF, Gas6 and Wnt5a were inactive or induced weak responses and were not considered for further experiments. We detected immediate and strong signaling response upon stimulation of cells with EGF, TGFα, IGF-1, HGF and TNFα (**Figure S1A**). We confirmed expression of the receptors of these ligands (EGFR, IGFR-1, Met and TNFRs), by measuring the abundance of mRNA in starved HeLa cells (**Figure 1D**). Quantitative analysis identified differences in the dynamics of signaling network activities between all five growth factors (**Figure 1E**). For example, in agreement with previous studies, TGFα induced much stronger signaling responses in comparison to EGF although both bind to the same receptor, EGFR (Ebner and Derynck, 1991; Francavilla et al., 2016; Scholler et al., 2017) (**Figure 1F**). Although TGFα induced a stronger response for most measured signaling events, stimulation with EGF or TGFα produced a similar dynamic trend of AKT and ERK kinase activity (**Figure 1F**). In turn, IGF-1 induced strongly sustained AKT activity, but led to a weak ERK activity followed by a slight inhibition after 10 min. In contrast, stimulation with HGF produced only relatively moderate transient activities of AKT and ERK while TNFα induced strong bi-phasic ERK activity and weak transient AKT activity. We used principal component analysis (PCA) to visualize the time-resolved differential activation of signaling pathways (**Figure 1G**). The PCA plot shows that time points belonging to the same treatment form well-separated clusters, indicating strong differences in global signaling dynamics. This separation was even more apparent in a t-SNE visualization when compared to PCA (**Figure S1B**). Importantly, these visualizations show that the effects of EGF treatment were highly similar to those of TGFα, indicating the strong similarity of the signaling program induced by growth factors binding to the same receptor, EGFR. Furthermore, we confirmed for the cells used here that extracellular cues such as GFs and cytokines are able to induce distinct dynamic signaling programs.

### Pairwise signalling interactions tune the network activity state

Following our hypothesis that two (or more) signaling cues can produce a different signal than a simple additive effect of individual growth factors, we reasoned that the signaling interaction between extracellular cues is not only dependent on the identity of the GFs but also on their relative concentration that is similar to EGF/IGF-1 interactions (Kuchenov et al., 2016). We therefore treated HeLa cells with three pairs of growth factors (TGFα/IGF-1, EGF/HGF and IGF-1/HGF) at various concentrations. The data on signalling network activity were combined with our previous EGF/IGF dataset obtained under the identical experimental setup in HeLa cells (Kuchenov et al., 2016). To assess cell-to-cell variation in the signalling response, we calculated the average coefficient of variation (CV) across all time points for each treatment/FRET biosensor pair. We found that 87% of FRET biosensor-treatment pairs had a CV <15% indicating strong consistency of single cell responses across different conditions (**Figure S2A**). Thus, the data confirmed the precision of our signaling network activity analysis. Principal component analysis showed a clear separation of the global signalling responses upon treatment with a different ratio of each pair of GFs (**Figure 2A-C**). Importantly, PCA illustrated that the shift of the cluster of time points belonging to the same treatment is not proportional to the concentration or to the ratio of GFs for all GF pairs tested. This result is in line with previous findings (Antebi et al., 2017; Kuchenov et al., 2016). Surprisingly, although EGF and TGFα bind to EGFR (Scholler et al., 2017) and induced similar signaling programs (**Figure 1G**), they exhibited different profiles of signaling interaction with IGF-1, suggesting that the interaction is also tuned by the identity of the cue with its unique signaling characteristics (**Figure 1E, S1B** and **2D**). The analysis of signaling interaction between the pairs EGF/TNFα and IGF-1/TNFα that activate distinct families of receptors, revealed a clear shift in PCA space in response to a combinatorial treatment, indicating similar mechanisms of signal integration (**Figure S2B**). As an independent confirmation, we observed that signaling signatures between EGF and IGF-1 occurred in human H838 non-small cell lung carcinoma cells (**Figure S2C**). Overall, these findings suggest that the mechanism of multiple cue signaling program tuning by down-stream interactions follows a general way of signal integration in living cells.

**Figure 2.**
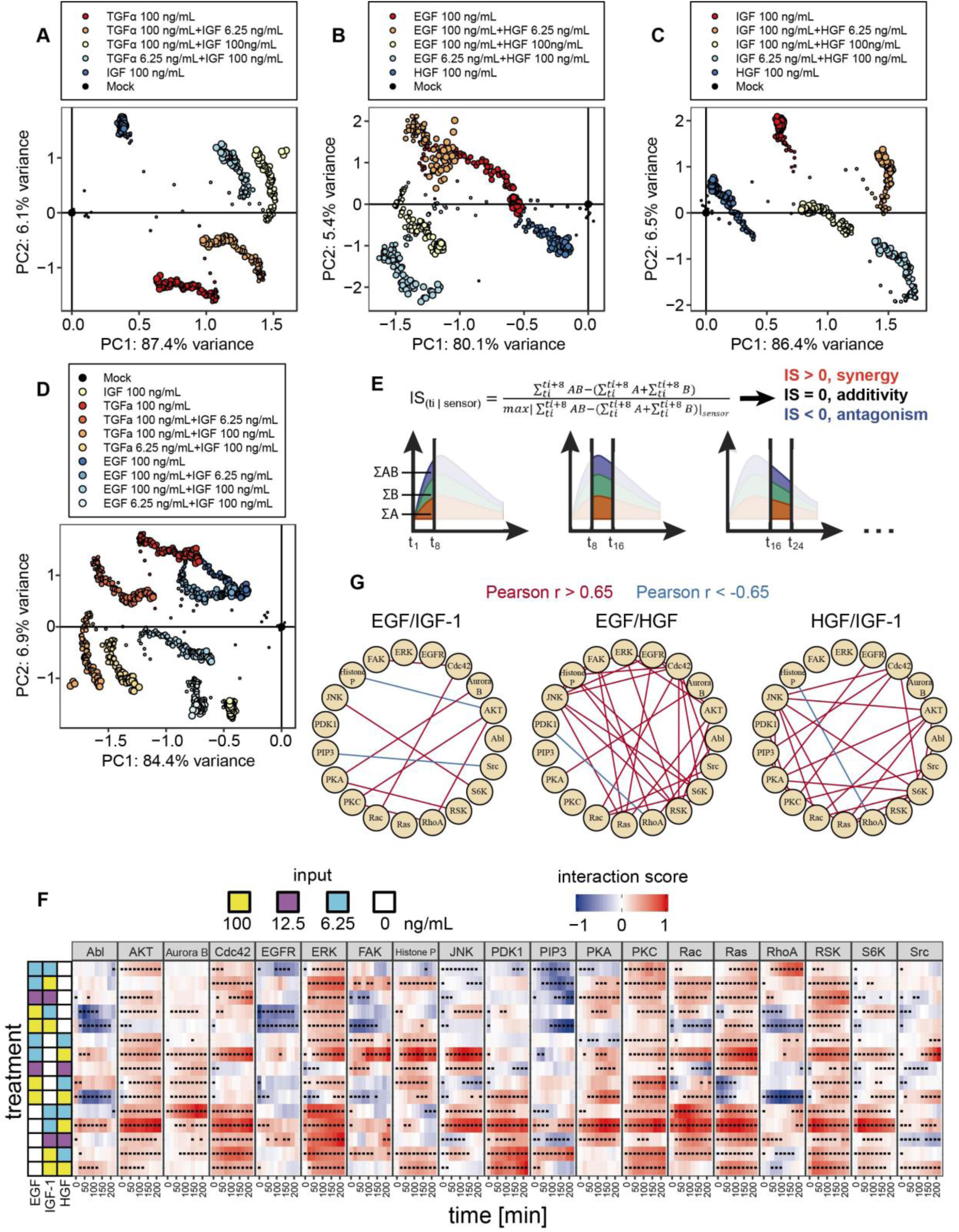
Induction of distinct dynamic signaling programs by a combinatorial GF code. (**A-C**) Global signaling response to the combinations of three ligand pairs projected into the first two principal components: (A) TGFα/IGF-1, (B) EGF/HGF, (C) HGF/IGF-1. Each dot represents a time point. (**D**) Comparison of global signaling response for TGFα/IGF-1 and EGF/IGF-1 using PCA. Each dot represents a single time point. (**E**) Calculation of the interaction score (IS) between growth factors. (**F**) The dynamic interaction map reflects antagonism, additivity, and synergy over time in response to stimulation with a pair of GFs. Signaling interaction is quantified and classified as depicted in (E). All significant IS values (adjusted p-value < 0.05, Student’s two sample t-test) are marked with a black dot. (**G**) Graph representations of strong Pearson correlations between interaction profile of signaling events for EGF/IGF-1, EGF/HGF and HGF/IGF-1. Red and blue edges represent positive (r > 0.65 with adjusted p value < 0.005, Student’s t test) and negative (r < −0.65 with adjusted p value < 0.005, Student’s t test) correlations, respectively.

To obtain new insights into the mechanisms underlying distinct signaling activity states in the presence of a pair of GFs, we classified down-stream signaling signatures into antagonistic, additive and synergistic, in a time-resolved manner. We defined an interaction score (IS) as the difference, in a specified time window, between the response to the combined stimuli A and B versus the sum of responses to the individual stimuli, subsequently divided by the maximum absolute IS value observed for each FRET sensor (**Figure 2E)**. The IS detects non-additive signaling interactions (synergy and antagonism) but remains largely unbiased with regard to the underlying molecular mechanisms. A positive IS value indicates synergistic behavior between stimulus A and B whereas a negative IS value suggests antagonistic or saturating behavior. At the same time, IS = 0 represents a purely additive response or the absence of a response. First, we confirmed that the most common interaction type was additive (IS is close to zero) indicating that our multiplicative normalization worked well over the full range of observed synergy and antagonism (**Figure S2D)**. This finding is also in line with data suggesting that additive interactions are dominant within signaling network(Natarajan et al., 2006). We then created a composite interaction map to visualize signaling interaction modes over time between each pair of growth factors (**Figure 2F** and **S2E**). Notably, the interaction mode between two extracellular cues was highly dynamic which is consistent with the PCA of the global signaling response (**Figure 2A-D** and **F**). All growth factor pairs displayed a distinct profile of IS, supporting previous observations by us and others (Borisov et al., 2009; Chatterjee et al., 2010; Kuchenov et al., 2016). Importantly, for each pair of GFs, the signaling interaction mode was dependent on the concentration and ratio of two stimuli explaining the non-proportional shift in the PCA space for each growth factor pair.

A major motivation for quantitatively measuring and systematically mapping of the IS between a pair of GFs is the possibility of identifying unanticipated cross-talk between them. We hypothesized that signaling molecules with highly correlated IS patterns tend to be directly or indirectly co-regulated under pairwise treatment. To assess GF pair-specific signaling interactions, we computed pairwise Pearson correlations of time-dependent IS patterns for all measured FRET biosensors for each GF pair: EGF/IGF-1, EGF/HGF and HGF/IGF-1 (**Figure S2F- S2H**). Next, we computed a correlation network to visualize the architecture of non-linear interactions for each growth factor pair (**Figure 2G**), where we defined the pair of sensors showing significant correlations (|r|>0.65, a Benjamini-Hochberg adjusted p-value < 0.005). This analysis revealed that different pairwise combinations of GFs resulted in a distinct network of synergistic and antagonistic cross-talk. Importantly, such analysis revealed strong positive correlation of the IS profile between PKA and RSK, specifically for the EGF/IGF-1 stimulation pair (**Figure 2G**). This is in line with the fact that RSK directly interacts with the catalytic domain of PKA upon stimulation with EGF (Gao and Patel, 2009). For all GF pairs tested, we detected positive IS profile correlation between JNK and S6K (**Figure 2G**). In support of this notion, it was previously shown that active JNK1 is able to directly phosphorylate S6K under various experimental settings (Martin et al., 2014; Miller et al., 2017; Zhang et al., 2013). These findings indicate that at least some of the detected correlations may be a result of direct interplay between two signaling molecules. Interestingly, the EGF/HGF pair showed a highly enriched positive correlation profile suggesting massive signaling interactions within the network (**Figure 2G**). In contrast, the EGF/IGF-1 pair exhibited a low-density correlation pattern (**Figure 2G**). Together, these results suggest that distinct concentration- and ratio-dependent combinatorial signaling interactions shape a dynamic signaling program and cause diverse signaling activity states.

### The impact of signalling interactions on the gene expression profile

Assessing the impact of a signaling activity state on a gene expression profile is fundamental in understanding the link between extracellular environment and the physiological state of the cell. We therefore investigated whether concentration- and ratio-dependent signaling programs of growth factor pairs were associated with changes in gene expression. We specifically focused on the EGF and IGF-1 since this pair induced distinct signaling programs (**Figure 1E-G**) and showed strong concentration- and ratio-dependent signaling interactions (**Figure 2D** and **F**). Using genome-wide RNA-seq, we quantified the mRNA abundance upon individual or combinatorial treatments in triplicates at 4 hours after stimulation (**Figure 3A**). We first confirmed that our RNA-seq analysis reproduced previously published data from HeLa cells treated with 20 ng/mL EGF (**Figure S3A**)(Amit et al., 2007). Biological replicates for different samples gave highly similar transcription profiles (**Figure S3B**), confirming the accuracy and precision of our RNA-seq analysis. As expected from differential pathway activation (**Figure 1E and F**), the differentially expressed genes (DEGs) induced by individual treatments varied considerably between EGF and IGF-1, especially at low GF doses (**Figure S3C**). We further globally examined the effect of individual and combinatorial treatments on the gene expression profile. PCA showed a pattern of distinct gene expression profiles that were similar to the pattern of the signaling network states (**Figure 3B and C**) indicating that concentration- and ratio-dependent signaling programs result in a distinct transcription output. The comparison of treated cells against control cells identified 9466 DEGs (DESeq2 method (Love et al., 2014), FDR= 0.05). Next, we directly compared DEGs between each condition (**Figure 3D** and **Table S1**). Although individual low concentration GF treatments resulted in a small number of DEGs, transcriptional changes in both directions were observed under each of the conditions (**Table S1**). Next, we were able to identify that each condition also regulated large unique gene sets, except for individual treatments of the lowest (6.25 ng/mL) concentration (**Table S1** and **Figure 3D**). Overall, these data suggested that concentration and ratio-dependent GF signaling programs result in complex changes of the gene expression profile in a non-additive manner.

**Figure 3.**
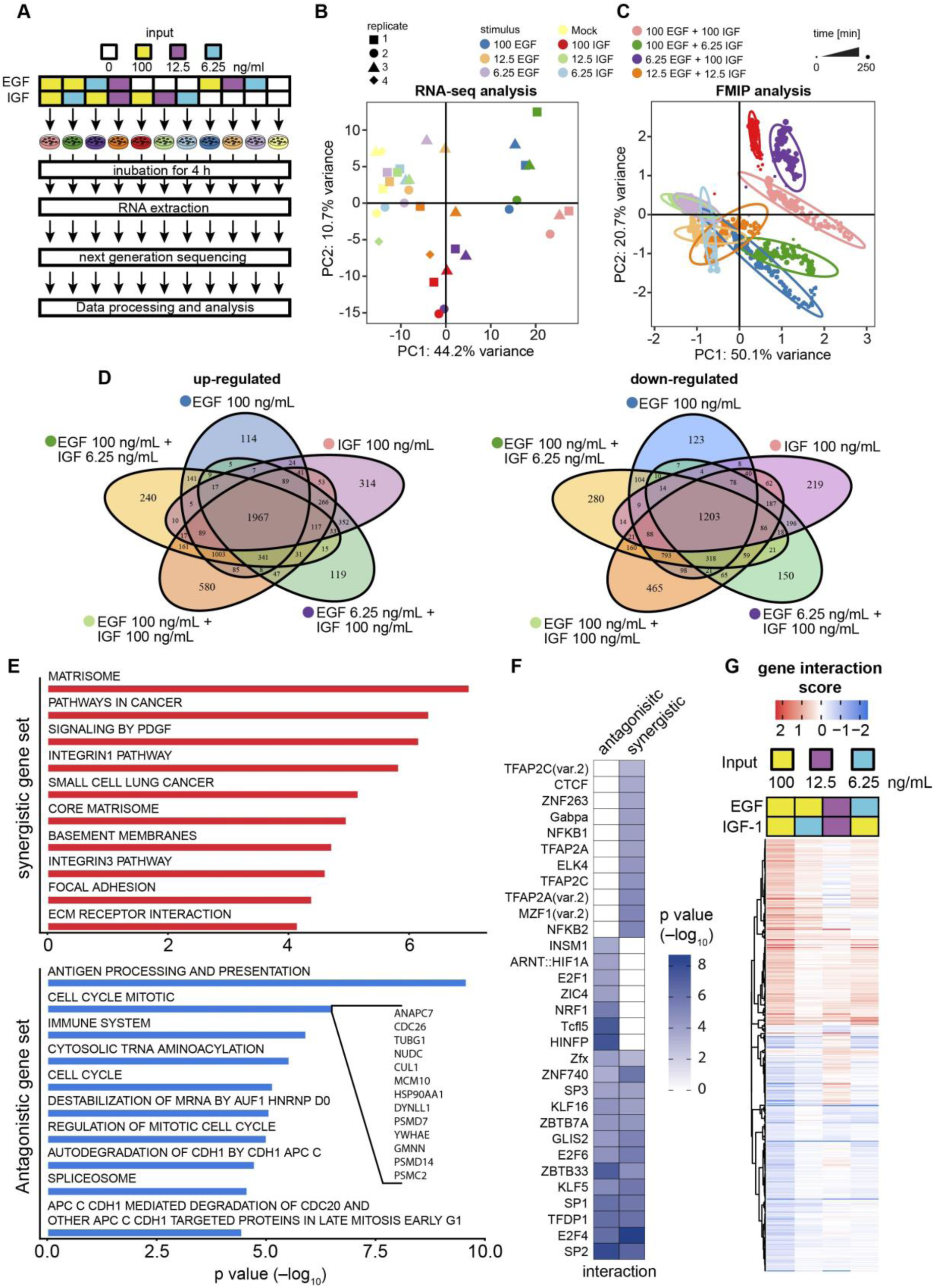
Combinatorial cytokine code governs gene expression profile through the signaling network activity state. (**A**) Experimental setup applied to determine combinatorial GF code-specific transcriptomes. HeLa cells were stimulated with various concentration combinations of EGF and IGF (individually or pairwise) for 4 h. Subsequently, cells were lysed, mRNA was extracted and subjected to RNA-seq analysis. (**B**) PCA based on 10000 genes of cells treated as indicated in (A) and measured by RNA-seq (n ≥ 3) (**C**) PCA based on FRET biosensor data measured by FMIP over 4 hours. Cells were stimulated as illustrated in (A). (**D**) Venn diagrams portraying significantly GF-induced/- suppressed genes and differential responses under the individual and pairwise treatments. (**E**) Gene set enrichment analysis of antagonistic or synergistic genes that responded to at least one pairwise treatment (adjusted p value < 0.05). (**F**) Heatmap shows known transcription factor (TF) binding motifs that were significantly enriched in each interaction response gene (IRG) group (with adjusted p value < 0.05 and |gene interaction score (GIS)| > 0.5). TFs with p value < 0.001 and RNA expression > 500 counts were defined as significantly enriched. (**G**) Heatmap representation for clustering of gene interaction scores for all synergistic or antagonistic genes that responded to at least one pairwise treatment (FDR – 0.1).

To gain insight into the molecular consequences that were triggered in response to the combinatorial EGF/IGF-1 treatment at the level of gene expression, we calculated a gene interaction score (GIS) to quantify non-linear transcription interactions induced by pairwise GF treatment. The antagonistic, additive and synergistic genes were then categorized in 9 groups according to the interaction profile similar to a recently published classification (**Figure S3D**) (Cappuccio et al., 2015). We found that the vast majority of interactions between EGF and IGF-1 were well approximated by an additive interaction, indicating that most gene expression changes are accurately predictable from individual treatments, which is in line with the observation that most genetic interactions are additive (Costanzo et al., 2016). Interestingly, for synergistic and antagonistic GISs the predominant influence on gene expression was buffering, where individual treatments induced changes in the same direction and the paired stimulation also resulted in an effect in that direction, but lower than the sum (**Figure S3E**). Further analysis of biological process enrichment for interaction response genes (IRGs) that responded synergistically or antagonistically to at least one pairwise treatment (adjusted p value < 0.05) shows overrepresentation of pathways linked to synergistic regulation of the extracellular matrix (ECM), focal adhesion, integrin, PDGF, and cancer related signaling (**Figure 3E)**. We observed a significant enrichment for antagonistic IRGs involved in the regulation of mitosis (cell cycle), splicing and mRNA destabilization (**Figure 3E)**.

We further searched for over-representation of transcription factor (TF) binding motifs in promoter regions of the 119 synergistic and 88 antagonistic genes (|GIS| >0.5 and a Benjamini-Hochberg adjusted p value <0.05) using Pscan (Zambelli et al., 2009) and the JASPAR non-redundant database (Khan et al., 2018). We identified a significant enrichment (p value <0.001) for sites binding to 31 TFs expressed in HeLa cells and classified them into three subgroups (**Figure 3F**): 11 TFs that exhibited the strongest enrichment in synergistic genes; 7 TFs that exhibited the strongest enrichment in antagonistic genes; and 13 shared TFs that were enriched in both. Importantly, among TFs enriched in antagonistic genes, we identified histone nuclear factor P (HiNF-P or MIZF), which interacts with methyl-CpG-binding protein-2 (MBD2) to promote DNA methylation and transcription repression (Sekimata and Homma, 2004). Among TFs enriched in synergistic genes, we found TFAP2A and SP1, both known for associated gene transcription activation in HeLa cells (Orso et al., 2010). Thus, we identified a putative number of TFs that tune gene expression in a combinatorial manner upon pairwise GF treatment.

Whereas synergistic and the antagonistic effects were observed under all tested concentrations and ratios of EGF and IGF-1, global analysis of IRGs suggested that the effects on individual genes were differed profoundly for each condition (**Figure S3F** and **3F**), mainly expressed as a fundamentally different interaction profile that was concentration and ratio dependent (**Figure 3F)**. Collectively, our data indicate that observed concentration- and ratio-dependent signaling states result in a distinct gene expression profile tuned by non-linear transcription interactions for regulating a diverse set of cellular processes and functions.

### Quasi-physiological environment shapes signaling and cellular response

The physiological levels of GFs are much lower than the concentrations that are typically applied in cellular signaling studies with serum-starved cells (Francavilla et al., 2016; Hill et al., 2016; Sigismund et al., 2005). Our experiments with low concentrations of growth factors revealed that pairwise treatments modulate the strength of signaling response depending on the identity, concentration and ratio of extracellular cues (**Figure 2F** and **S2D)**. Furthermore, gene expression analysis demonstrated the boost of total DEGs upon co-treatment with EGF and IGF-1 of low concentration (12.5 ng/ml) (**Figure 4A** and **S4A**) indicating strong interaction of EGF and IGF-1 at low concentrations on the level of gene expression. Therefore, we reasoned that in the presence of multiple extracellular cues at low concentrations typically provided by 10% fetal bovine serum (FBS) the signaling network state is adjusted similar to multiple GF additions (**Figure 2F**) and hence leads to altered sensitivity and response patterns. We therefore studied the dynamic signaling program upon stimulation with EGF at low and high concentrations in overnight serum starved and non-starved HeLa cells. As expected, the EGF-induced signaling program was altered in the presence of 10% FBS (**Figure 4B**). Notably, the signaling response after a low concentration of EGF (6.25 ng/ml) was much stronger in the presence of FBS compared to serum starved HeLa cells. (**Figure 4B** and **C**). For example, we observed a stronger response of Src, ERK, and S6K in non-starved cells (**Figure 4C**). Immunoblot analysis qualitatively confirmed that the presence of 10% FBS elevated EGF-induced pS6 levels, confirming the FRET data (**Figure 4D** and **S4B**). Next, we analyzed the global signaling response by PCA in serum starved and non-starved cells. We observed a clear separation between serum starved cells treated with a range of EGF concentrations and non-starved cells, indicating that the presence of 10% FBS tunes the cells to respond specifically to the EGF stimulation (**Figure 4E**). In contrast, PCA showed that the presence of FBS resulted in a relatively modest effect on the signaling responses when cells were stimulated with a high concentration of EGF which is in line with a previous report (Lun et al., 2017) (**Figure 4E**). Together, these data indicate that the presence of multiple extracellular cues considerably tunes the signaling dynamic program in response to the GF stimulation of low concentration. This has major implications for comparing results of starved and non-starved cells in cell biology experiments.

**Figure 4.**
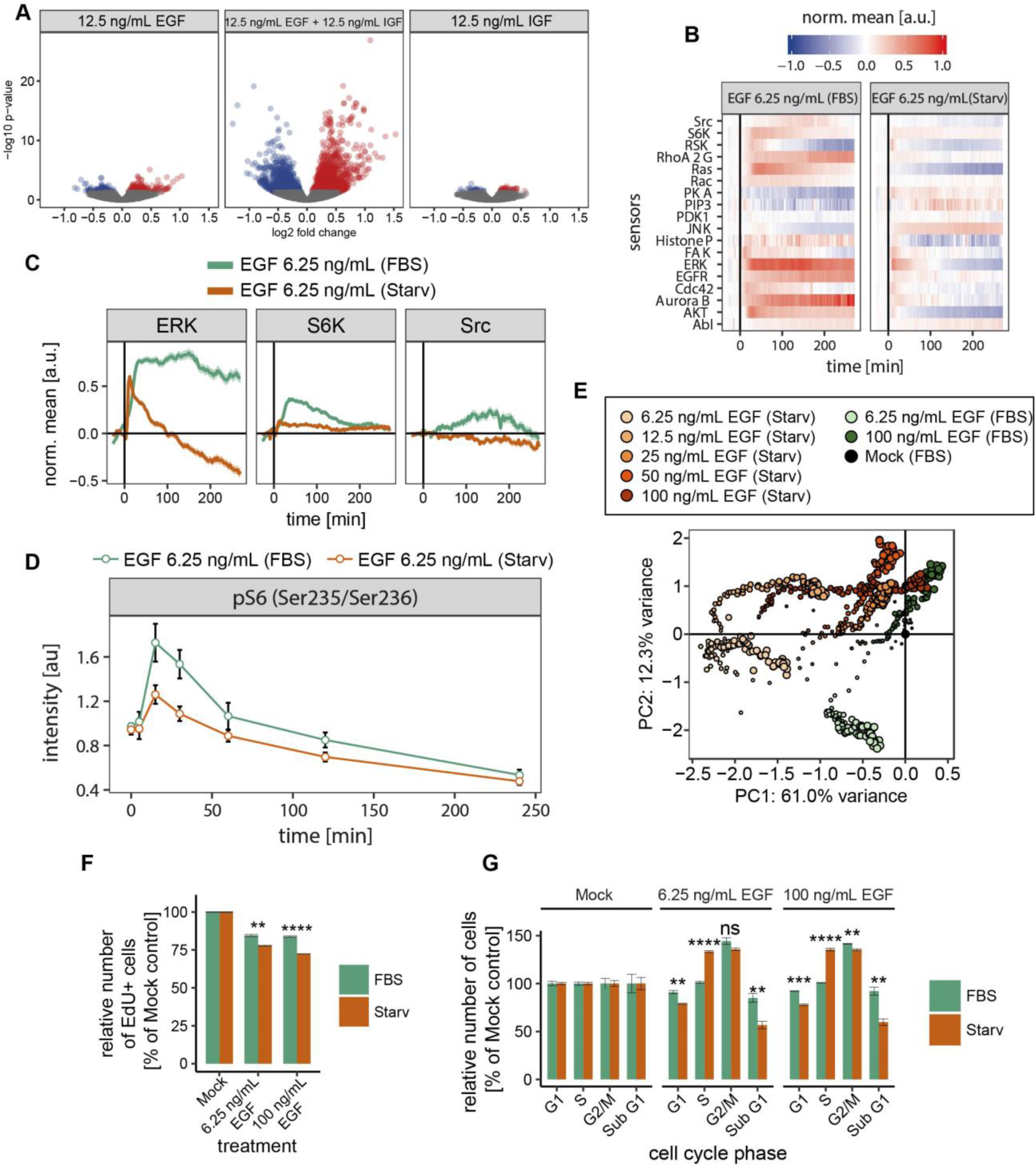
Quasi-physiological condition potentiates and shapes signaling response. (**A**) Volcano plots comparing cells treated with 12.5 ng/mL EGF, 12.5 ng/mL IGF-1 and 12.5 ng/mL EGF + 12.5 ng/mL IGF-1. Data represents three independent experiments. Blue and red dots indicate suppressed and induced genes with adjusted p value < 0.05, measured by DESeq2, respectively. (**B**) Quantification of the dynamic program of signaling activity after exposure to 6.25 ng/mL EGF in absence or presence of 10% FBS. (**C**) Examples of potentiation of signaling events in the presence of 10% FBS from (B). (**D**) Phosphorylation of S6 detected by quantitative immunoblotting. HeLa cells were maintained in either growth factor depleted or 10% FBS supplemented medium and stimulated with 6.25 ng/mL EGF. Error bars represent ± SEM. (**E**) Global signaling response to a range of EGF concentrations projected into the first two principle components. (**F**) Relative 5-bromodeoxyuridine (EdU) incorporation for cells kept in 10% FBS or serum starved cells. Cells were left unstimulated (mock) or stimulated with 6.25 ng\mL or 100 ng\mL EGF. Error bars represent ± SEM. (**G**) Comparison of the relative change of each cell cycle phase (G1, S and G2/M) between cells kept in 10% FBS supplemented or serum deprived medium. Cells were left unstimulated or stimulated with 6.25 ng/mL or 100 ng/mL EGF. Error bars represent ± SEM. Significance was assessed by Student’s t test. ns: p > 0.05, *: p ≤ 0.05, **: p ≤ 0.01, ***: p ≤ 0.001, ****: p ≤ 0.0001.

To determine the functional relevance of signaling difference induced by the presence of 10% FBS, we quantified proliferation and determined cell cycle parameters assayed by 5-ethynyl-2-deoxyuridine (EdU) incorporation and 4,6-diamidino-2-phenylindole (DAPI) staining, respectively. As expected, serum starvation resulted in a reduced basal proliferation as well as increased fraction of cells in G1 phase (**Figure S4C**). Stimulation with EGF resulted in the inhibition of cell proliferation as well as an increase in the percentage of cells in G2/M phase regardless of the starvation status, a well-known functions of EGF on cancer cell lines (Gill and Lazar, 1981; Kinzel et al., 1990; MacLeod et al., 1986) (**Figure 4F-G** and **S4C**). This finding is also in line with the fact that mitotic cell cycle genes were highly enriched in the antagonistic gene set (**Figure 3E**). We further quantified that HeLa cells subjected to starvation and treated with EGF demonstrated a stronger decrease in proliferation in comparison to cells treated with EGF in the presence of 10 % FBS (**Figure 4F**). In contrast, EGF treatment resulted in a significant increase in the relative number of starved cells in S phase than cells kept in full media. The latter showed a strong decrease in the number of cells in G1 phase (**Figure 4G**). Collectively, these data suggest that the molecular composition of FBS tunes the signaling network state: 1) to achieve higher signaling responsiveness of the cells and 2) to shape the EGF-induced signaling program and physiological response under physiologically relevant concentration.

### Quasi-physiological environment tunes the basal signaling network activity state

Recent work suggested that the processing of information from multiple cues might occur at the receptor level (Antebi et al., 2017). To address this possibility, we quantified total and plasma membrane EGFR expression by immunoblot analysis and immunostaining, respectively. Importantly, although we observed a slight increase in the expression of plasma membrane EGFR upon starvation (**Figure 5A** and **5B**), we did not detect a significant difference in total EGFR as well as in ABL and SRC expression between cells kept in full media and serum starved cells (**Figure S5A, S5B and S5C**). Our findings indicate that tuning the signaling program by the extracellular environment occurs within the intracellular signaling network, i.e. by adaptation of the basal activity of downstream components. These findings encouraged us to investigate the mechanism of tuning the signaling response in the presence of multiple extracellular cues of low concentrations. First, we compared the basal activity state of the signaling network in the presence or absence of 10% FBS. Interestingly, basal signaling profiling revealed that all signaling events in our data set exhibited a high variability at the single cell level. As expected, we found that basal patterns were classified into three groups: signaling events either showed a decrease, an increase or no difference in basal activity upon starvation (**Figure 5C**). These findings are in line with a previous study (Pirkmajer and Chibalin, 2011) and further support our hypothesis that a signaling network is tuned under quasi-physiological conditions to achieve a specific basal activity state.

**Figure 5.**
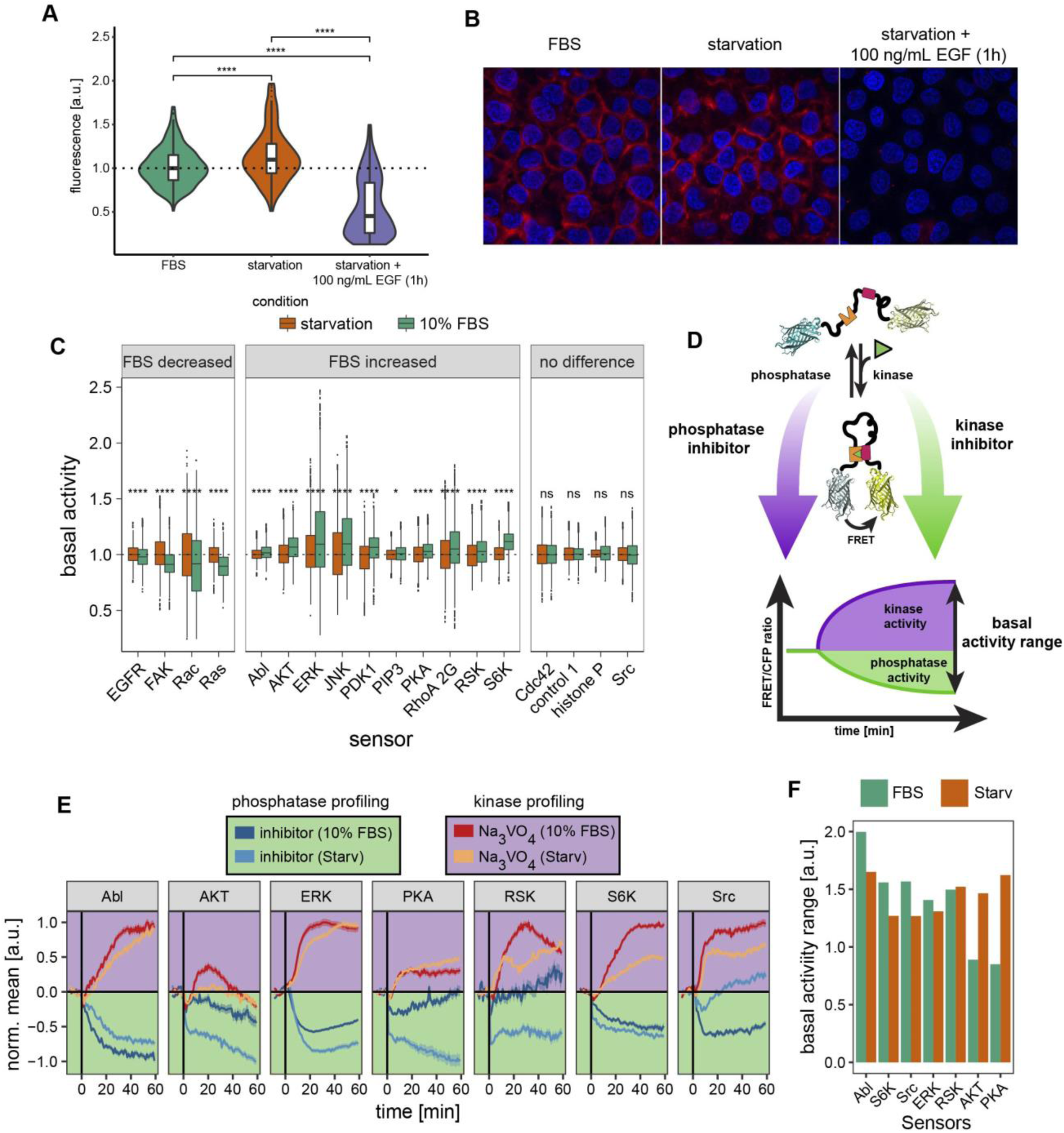
Complex environment fine-tunes steady-state signaling. (**A**) Quantification of the EGFR surface abundance by staining with an antibody against the extracellular EGFR epitope. The fluorescent intensity (y axis) under various conditions (x axis) is depicted. Fluorescence intensities of the receptor were normalized to those in FBS. (**B**) Representative images from (A). (**C**) Basal activity profile in serum starved and non-starved cells acquired with FRET biosensors. (**D**) Illustration of the experimental setup to study kinase and phosphatase basal activity. (**E**) Quantification of kinase and phosphatase basal activity upon tyrosine phosphatase or kinase inhibition in serum starved and non-starved cells. Cells were treated with 10 µM multi-AGC kinase inhibitor (AT13148; AKT, PKA, RSK, S6K biosensors), 10 µM MEK inhibitor (AZD6240; ERK biosensor), 10 µM Src/Abl dual inhibitor (SKI-606; RSK and ABL biosensor) or 100 µM protein-tyrosine phosphatase inhibitor (activated Na_3_VO_4_; all biosensors). Mean ± SEM is shown. P values are derived from Wilcoxon test. ns: p > 0.05, *: p ≤ 0.05, **: p ≤ 0.01, ***: p ≤ 0.001, ****: p ≤ 0.0001. (**F**) Quantification of basal activity range from (E).

Next, to investigate how the interplay between multiple extracellular cues might cooperate to mediate distinct signaling states, we took advantage of those FRET biosensors that are able to monitor the balance between kinase and phosphatase activity (Sample et al., 2014). We applied inhibitors to dissect the underlying mechanisms of tuning basal signaling activity by using FRET sensors as readout of kinase and phosphatase activity (**Figure 5D**). Upon treatment of serum starved cells with activated sodium orthovanadate (Na_3_VO_4_), a general protein-tyrosine phosphatase inhibitor (Huyer et al., 1997), we observed that the majority of biosensors including EGFR, ERK, RSK, S6K, SRC, and FAK exhibited elevated FRET ratios supporting the fact that kinase signaling is active in the absence of ligand, but is continually repressed by phosphatase activity in living cells (Reddy et al., 2016) (**Figure S5D**). Importantly, we did not observe AKT kinase activity upon starvation. In addition to the changes in kinase biosensors, we detected elevated activity of RhoA and Cdc42 GTPases in the presence of orthovanadate (**Figure S5D**). We identified that AKT, RSK, S6K, and SRC phosphorylated FRET biosensors more efficiently in the presence of 10% FBS compared to serum starved cells (**Figure 5E**). In contrast, JNK and PKA were slightly less active in the presence of serum. Together, these results suggest that the steady-state activity of tyrosine kinases is under a complex environmental control.

We further profiled basal phosphatase activity that is regulating the phosphorylation level of the SRC, AKT, S6K, RSK, ERK, ABL, and PKA FRET biosensors by inhibiting kinase activities. The SRC/ABL dual inhibitor (SKI-606) induced a stronger dephosphorylation level of the SRC and ABL biosensors in the presence then in the absence 10% FBS, suggesting a higher activity of phosphatases compared to starved cells (**Figure 5E**). In contrast, the treatment with the multi-AGC kinase inhibitor AT13148 as well as the MEK inhibitor AZD6240 resulted in a stronger dephosphorylation of the AKT, S6K, RSK, PKA, and ERK FRET biosensors upon serum starvation than in the presence of 10% FBS (**Figure 5E**). We further quantified the basal activity range (**Figure 5D**) in serum starved and non-starved cells. Notably, we detected a strong increase in basal activity range by the ABL, S6K, and SRC FRET biosensors in the presence of 10% FBS. In contrast, the AKT and PKA FRET biosensors reported an increased basal activity range upon starvation. Collectively, these results demonstrate that an extracellular environment regulates the basal phospho-turnover rate and the basal activity range of the signaling network. More broadly, our results reveal that information from a signal-rich environment can be encoded by the basal phosphorylation status and phospho-turnover to fine-tune a response to a newly incoming cue.

## DISCUSSION

The current prevailing view is that aberrant signaling programs under pathological condition are predominantly induced by mutations or large changes in protein abundance (Adlung et al., 2017; Hill et al., 2016; Li et al., 2017; Lun et al., 2017). However, other recent studies show that the extracellular microenvironment dramatically changes in diseased conditions compared to normal (Rezza et al., 2016; Sever and Brugge, 2015). Such changes in the microenvironment through autocrine and paracrine GF/cytokine signaling may result in altered signaling dynamics (Knapp et al., 2017). Here we used a FRET biosensor-based multiparameter imaging platform (FMIP) (Kuchenov et al., 2016) to investigate how cells integrate information from more than one input cue. FRET sensors intrinsically monitor enzymatic activity continuously over time in single cells. Therefore, FMIP reports on the functional state of a signaling network over extended periods of time, here for four to five hours. This is an essential advantage over immunoblotting blotting based approaches that report on the posttranslational modification status of proteins only at the bulk level and at a limited number of time points. The FMIP has proven to be an effective research tool to investigate signal integration from multiple distinct inputs. Our FMIP assay reveals a broad landscape of signaling interactions such as antagonism, additivity, and synergy after stimulation of a group of distinct receptors (**Figure 2F)**. The downstream signaling signatures resulted in a dynamic and unique signaling program that is specific for a given combinatorial input. Such stimulation scheme serves as a simplified model for the complex cues provided by the extracellular microenvironment. We demonstrated at the systems-level that the time-dependent signaling signatures (antagonism, additivity, and synergy) were distinct for each condition and depended on the concentrations as well as ratio of the ligands. Importantly, we showed that the signaling signature are arranged in a complex pattern of signaling cross-talk between components of distinct pathways (**Figure 2G)**. Thus, although we cannot rule out the contribution of receptor trafficking in the regulation of signaling network activity, our data on pairwise stimulation clearly indicate that ratio sensing of GFs is exerted by the signaling network of the respective receptors.

The identification of signaling interaction modes regulating distinct dynamic signaling responses raises a question: are the interaction modes biologically relevant? Our RNA-seq analysis revealed that each signaling network state resulted in a distinct gene expression profile as well as in a distinct interaction profile at the level of transcription (**Figure 3B-D** and **G**). Our observations on HeLa cells correlate well with previous studies obtained from several cell types of distinct species where strong combinatorial interactions at the level of transcription have been observed for growth factors and cytokines (Ammeux et al., 2016; Cappuccio et al., 2015; Martin et al., 2009). Although we identified a large number of concentration- and ratio-dependent interactions at the level of gene expression upon pairwise treatment with EGF and IGF, many of them appeared weak under low concentrations. Thus, the integration from a much more complex combinatorial treatment of low concentrations relevant to physiological conditions will produce a biologically meaningful effect on gene expression.

We paid particular attention to kinase and phosphatase activities, as their balance plays a critical role in signal transduction (Day et al., 2016). Our experiments using kinase and phosphatase inhibitors while monitoring kinase and phosphatase activities over time suggested that the balance between protein tyrosine phosphatase and kinase activity may be of greater significance in cellular adaptation to the changing environment than so far realized. We find evidence for the massive regulation of the steady-state turnover rate of protein phosphorylation and basal phosphorylation status under signal-rich (10% FBS) and serum-deprived conditions (**Figure 5C, E** and **F**). It was proposed that high rates of the activation/deactivation cycle may increase responsiveness and allow kinetic proofreading (Kleiman et al., 2011; Lemmon et al., 2016; Swain and Siggia, 2002). The differing patterns of enzymatic activity in response to a low concentration of EGF (**Figure 4B-E**) may therefore stem from distinct turnover rate of phosphorylation sites. Thus, our current findings provide support that similar mechanisms may underlie increased sensitivity and proof reading under exposure to multiple cues of low concentrations, as is typical for many tissues and microenvironments.

Based on our data, we propose that the concentration- and ratio-dependent signaling interactions of two or more cues are crucial for signal transduction under physiological conditions (**Figure 6**). Through such non-additive interactions, the extracellular environment that is defined by the identity of its cues, their concentrations and ratios, tunes the dynamic signaling program in response to a newly appearing cue. In support of this hypothesis, we showed that under quasi-physiological conditions, in the presence of multiple signaling cues at low concentrations (in the presence of serum), the cellular signaling network is pre-activated and tuned to achieve specific strong responses to low concentrations of ligands (**Figure 4B-E**). Although, it has long been known that serum is able to potentiate EGF- and IGF-1-induced responses (Draghi et al., 1980; Mandl et al., 2002), the molecular mechanisms of signaling network tuning and pre-activation by multiple extracellular cues in the order of seconds/minutes have not been explicitly identified yet. Undoubtedly, the signaling program is determined by kinetic parameters of the downstream reactions (network activity), network component abundance and their interactions (network topology) (Kolch et al., 2015; Purvis and Lahav, 2013). A long-standing question in the signaling field asks how the complex extracellular environment tunes such signaling parameters. In our study, we systematically illustrated that the underlying mechanisms of basal signaling activity state governing by multi-cue environment are linked to two coupled but conceptually distinct regulatory levels that work at different time scales: (1) basal protein abundance and network topology are controlled by gene expression and/or protein degradation that works on the order of minutes/hours (**Figure 3**) and (2) basal phosphorylation (**Figure 5C**), network topology (**Figure 2G**), phospho-turnover (**Figure 5E**), and basal activity range (**Figure 5F**) are regulated through signaling crosstalk emerging on the order of seconds/minutes. These two tightly linked regulatory levels allow the signaling network state to encode information from a combination of past experience and newly incoming signals.

**Figure 6.**
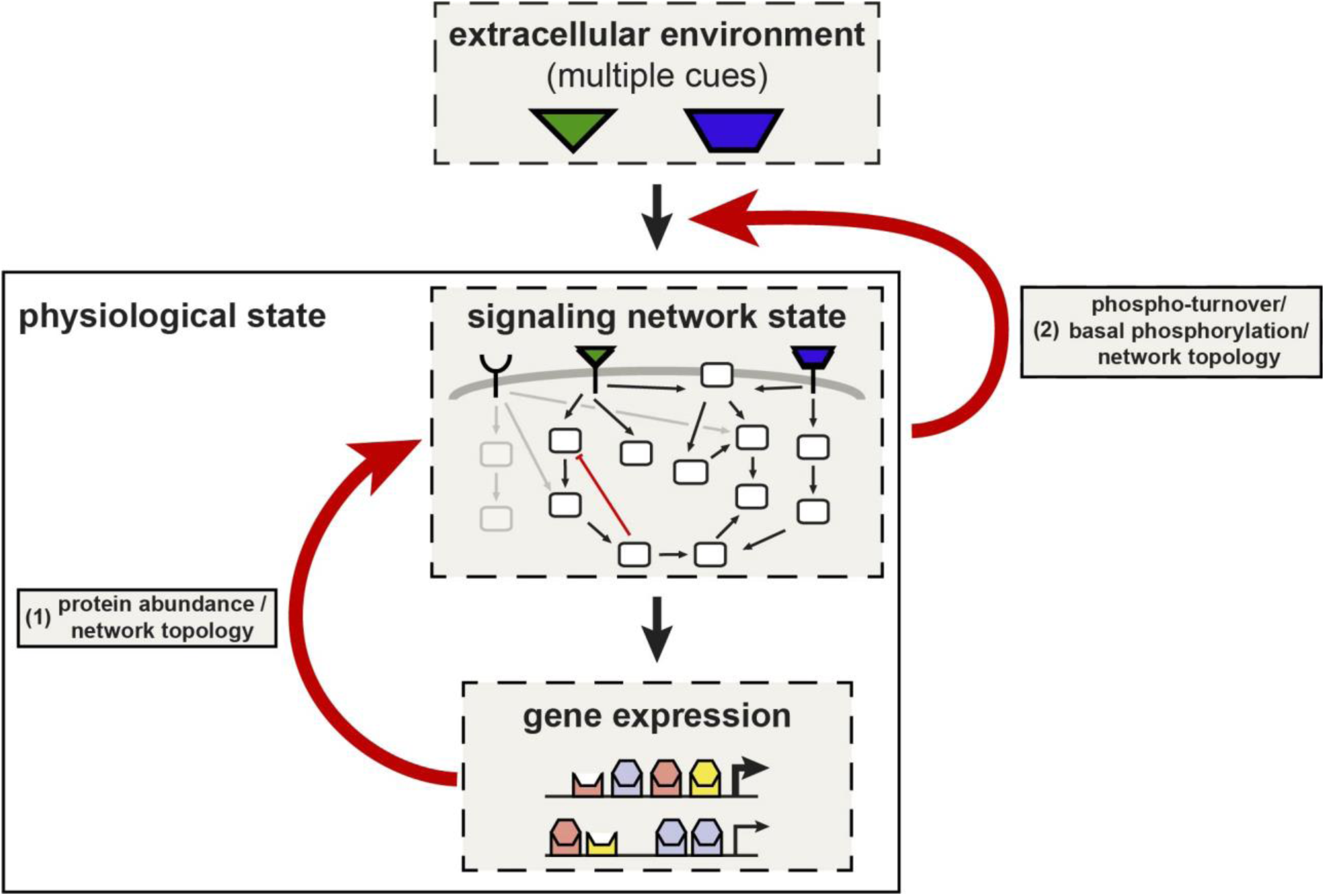
Activity-based network model for signal processing in the presence of multiple extracellular cues. Our study supports a feedback control scheme for extracellular information processing. First, basal protein abundance and network topology are dynamically controlled by gene expression and/or protein degradation, both well-described mechanisms. Second, basal phosphorylation, phospho-turnover and network topology are combinatorically regulated by the current extracellular environment through down-stream signaling interactions (or cross-talk). Both feedback loops affect the basal signaling activity state that in turn shapes the signaling program in response to a newly appearing cue through additivity, synergy and antagonism.

The activity-based model we propose could have far-reaching consequences for understanding how cancer cells adapt to the therapeutic inhibition of activated receptors or pathways. It is generally well established that the activation or amplification of the c-Met receptor tyrosine kinase is able to induce drug resistance in EGFR family mutant cancers (Bardelli et al., 2013; Minuti et al., 2012; Turke et al., 2010; Watson et al., 2018). However, the driving mechanisms leading to resistance are unclear. Intriguingly, we showed that low concentrations of HGF exhibited a strong synergistic effect on global EGF and IGF-1 signaling (**Figure 2G**). Thus, it is tempting to speculate that potentiation of the signaling network activity might be a driving force in the selection of beneficial subclones in response to therapeutic pressure.

In conclusion, our data determine a comprehensive activity-based network model of signal processing in the presence of multiple extracellular cues. Further studies based on this conceptual model will likely unveil novel mechanisms of cell adaptation to environmental changes including drug action and of the evolution of drug resistance.

## SUPPLEMENTARY INFORMATION

Supplementary information includes five figures and one table and can be found with this article online.

## ACKNOWLEDGMENTS

We thank Pedro Beltrao for critically reading the manuscript and all members of the Schultz lab for helpful comments. We are grateful to Rainer Pepperkok and Ekaterina Korotkevich for useful discussions. We are also grateful to the ALMF team at EMBL for excellent technical support and B. Neumann for help with microarray spotting. We thank the EMBL Genomics Core Facility for performing the next-generation sequencing analysis. This work was supported by the EMBL and the DFG (SFB 1129) (to C.S.); the Joachim Herz Stiftung (to D.K.); German Federal Ministry of Education and Research (BMBF) through the LungSysII consortium and the German Center for Lung Research (DZL) (to U.K. and C.S.); and the European Commission (SOUND research project) (to W.H.)

## AUTHOR CONTRIBUTION

C.S. and D.K. designed the study, D.K. performed most of the experiments and statistical analysis, F.Z., D.K., and W.H. performed statistical analysis of the RNA-seq data, F.S. performed immunoblot analysis. D.K. and C.S. wrote the manuscript with suggestions from all authors. U.K. contributed critically on levels of the study. C.S. and U.K. provided material support. The project was led by D.K. and supervised by C.S.

## METHODS

### Contact printing

The fabrication of microarrays was performed as described previously in (Kuchenov et al., 2016). Plasmids for reverse transfection were diluted to concentration of 1 mg/mL. The transfection mixture was prepared by mixing 9 µl of a 0.4 M sucrose solution in DMEM, 9 µl of DNA and 33µL of lipofectamine 2000 mixed in a 96-well plate and incubated for 20 min at room temperature. Subsequently, 21.75 µL of 0.29% gelatin solution in water was added to the transfection mixture. The transfection cocktail was distributed in 384-well plates (24 µL per well). The plates were stored at −20°C. In order to print Labteks the 384-well plate was thawed at room temperature and centrifuged briefly up to 54 g to straighten the surface of the samples. Afterwards, residual bubbles were removed by a 10 µL tip and placed immediately in the contact printer. Before printing, LabTek dishes were washed with 70% ethanol increasing hydrophobicity of the LabTek surface and, accordingly, improving the shape of the spots. 1-well LabTek dishes were printed with a “ChipWriter” contact printer equipped with solid pins. Using PTS 600 pins, the diameter of printed spots was about 600 µm and the spot-to-spot distance was 1125 µm. Printed 1-well LabTek dishes were stored at room temperature in a gel drying box in the presence of drying pearls (Sigma).

### Live cell imaging

HeLa (650,000) or H838 (800,000) cells were transiently transfected by seeding on printed LabTeks. The seeding was performed when cells were cultured to 50%-60% confluence. After maintaining HeLa and H838 cells in an incubator for 24 or 48 hr, respectively, the media were changed to starvation media 12–17 hr prior to imaging. During imaging, cells were maintained in imaging medium (minimum essential medium supplemented with 100 U/mL of penicillin, 100 mg/mL of streptomycin, and 30 mM of HEPES) at 37 C without CO2. To assist cellular segmentation cells were preincubated with 7.5 nM DRAQ5 for 30 min (Cell Signaling Technology).

Time-lapse imaging was performed on an Olympus IX83 microscope equipped with a Hamamatsu ImagEM CCD camera and an environmental control unit incubation chamber using 20x 0.70 numerical aperture (NA) or 10x 0.40 NA. The microscope was equipped with 436/20 excitation filter, a CFP/yellow fluorescent protein (YFP) dualband beam splitter (51017bs; Chroma), and two emission filters (470/30 for CFP and 535/50 for YFP) that were controlled by a filter wheel. Image acquisition was controlled by the Olympus IX83 microscope software – xCELLence (Olympus) or cellSens (Olympus). Fluorescence images were acquired with an exposure time of 200 ms.

### RNA extraction and sequencing

Hela cells were seeded in 60 mm dishes pre-coated with gelatin to reproduce conditions similar to microarrays. Prior to stimulation cells were starved in the absence of serum overnight. Afterwards, cells were incubated with GFs for four hours. We lysed cells by the direct addition of 3 ml of TRIzol (Life Technologies) after media removal according to the manufacturer’s instructions. For RNA extraction lysates were treated with 0.6 ml chloroform (Sigma-Aldrich) for 3 min and centrifuged at 12,500 g for 15 min at 4 °C. Aqueous supernatant was collected and diluted 1:1 with 70% ethanol. Total RNA was extracted from solution using RNeasy Mini Kit (Qiagen), following the manufacturer’s instructions and quantified using the NanoDrop spectrophotometer. RNA was used with A(260/280) nm ≥ 1.8 and A(260/230) nm ≥ 2.0. RNA quality was assessed using RNA 6000 Nano chips on the Agilent 2100 Bioanalyzer. The library preparation, RNA sequencing and reads alignment were performed by a Genomics Core Facility at EMBL. Sequencing was performed on Illumina NextSeq-500 instruments.

### Immunofluorescence

HeLa cells were seeded in a 12-well glass-bottom dish (Ibidi, 81201) and maintained in CO2 incubator for 24h. Afterwards, the medium was changed to full or FBS-free medium. After, 17h of incubation, cells were fixed with 4% paraformaldehyde (PFA; Electron Microscopy Sciences, 19208) in PBS for 10 min at room temperature and washed with PBS. Cells were blocked in 1% (w/v in PBS) BSA for 1 h at room temperature. After removal of the blocking solution (without any wash), cells were incubated with the anti-EGFR primary antibody (Santa Cruz Biotechnology, sc-101, 1:200) diluted in 1% (w/v, in PBS) BSA at 4 °C overnight. After washing with 1% (w/v, in PBS) BSA, cells were incubated with goat anti-mouse Alexa Fluor 546 (Life Technologies, A-11031) at room temperature for 1 hour. Cells were then washed with PBS at room temperature and finally rinsed with distilled water. After drying, coverslips were covered with SlowFade Diamond antifade mountant with DAPI (ThermoFisher Scientific, S36968). Microscopy images were captured at room temperature using a confocal laser scanning microscope (Zeiss LSM780) with a 63× oil objective and fully opened pin hole. Settings were as follows: DAPI-channel: 405-nm excitation (ex), 409- to 475-nm emission (em); red channel: 561-nm ex, 569- to 655-nm em. Images were further processed using Fiji software (fiji.sc/) with the in-house developed macro.

### Quantitative immunoblot analysis

For the analysis of phosphorylated and total S6, total EGFR and total Abl, 6·10^5^ HeLa cells were seeded one day in advance in a 4 cm cell culture dish (TPP) and washed three times with DMEM without additives and then incubated for 16 hours in growth factor depletion medium or growth medium containing 10 % FBS. The cells were stimulated with the indicated doses of EGF (GF144, Millpore) for the given timespan. The cells were lysed and processed as previously described (Merkle et al., 2016). Cell lysates were separated by SDS-PAGE and transferred to a PVDF membrane (0.45 μm pore size, Immobilon, Millipore). Antibodies against phosphorylated S6 (#2211), total EGFR (#4267), total Src (#2108) and total Abl (#2862) were obtained from Cell Signaling. Antibodies for loading control were purchased from Santa Cruz (anti-vinculin, sc-55465) and Sigma (anti-actin, A5441). Secondary antibodies coupled to hrp were obtained by Dianova. Luminescence was detected by ImageQuant LAS4000 (GE Healthcare) and signal intensity was quantified by ImageQuant TL software (Version 7.0, GE Healthcare).

### EdU incorporation assay

HeLa Kyoto cells were seeded into two 15 cm dishes at 20% confluency in DMEM supplemented with 10% FBS (DMEM + 10% FBS). 24 hours later, one dish (A) was washed once with DMEM only, and then serum-starved in DMEM only for 16 hours. The other dish (B) was washed once with DMEM + 10% FBS, and then cultured in DMEM + 10% FBS for 16 hours. At the end of the 16-hour incubation, cells on both dishes were detached using TrypLE Select (1X) and collected by centrifugation. Cells from A were washed once with DMEM only, and then re-suspended in DMEM only at 2 × 10^5^ cells/mL density. Cells from B were washed once in DMEM supplemented with 10% FBS, and then re-suspended in DMEM + 10% FBS at 2 × 10^5^ cells/mL density. 1 mL of suspensions from A and B were added to 35 mm dishes that already contained 1 mL of 2X concentration of the growth factors prepared in DMEM only and DMEM + 10% FBS, respectively. 24 hours later, 20 µL of EdU (1 mM) in PBS was added into each dish (final concentration 10 µM) for a 2-hour period. One dish was not treated with EdU and used as negative control. At the end of EdU labelling, cells were washed with PBS, detached from the dishes using TrypLE Select (1X), washed by centrifugation with PBS, and then fixed in 70% ethanol at +4 °C overnight. The next day the ethanol was removed, and then a copper-catalyzed click reaction was performed using Alexa 488 - picolyl azide supplied in Click-iT® Plus EdU Alexa Fluor® 488 Flow Cytometry Assay Kit from Thermo Fisher (#C10632). Cells were counterstained using DAPI (10 µg/mL final conc.), and finally analyzed for proliferation and cell cycle by BD LSRFortessa (BD Biosciences) cell analyzer.

## QUANTIFICATION AND STATISTICAL ANALYSIS

### Image analysis

The single image of a nuclei channel and the time-lapse series of CFP and FRET channels acquired from different positions were exported as separate TIFF files. The images were analyzed with FIJI (Schindelin et al., 2012) and FluoQ (Stein et al., 2013). The FIJI-based macro developed in-house was used to preprocess and multiply nuclei image and subsequently concatenate all channels together. This macro produced a single tiff file containing three channels: binary mask of nuclei (repeated 100 times), FRET and CFP channels obtained from the same position. The file containing three channels was further analyzed with the FIJI-based macro FluoQ. Although cells express different amount of FRET biosensors we used image analysis pipeline that automatically account for a low expressing cells that are close to background cellular fluorescence. In order to subtract background we used a histogram-based “Triangle” algorithm to calculate the mean of the thresholded background that was subsequently subtracted from each pixel. Next, image smoothing with a median filter (radius size = 2) was applied and the images were transformed to a 32-bit float. Cells were automatically segmented by using binary mask created by Huang’s fuzzy thresholding method. The signal intensity that was equal or close to intensity of the background was set as NaN value due to Huang’s fuzzy thresholding. This image analysis pipeline automatically accounts for a very low expressing cells and avoided erroneous FRET ratios. The nuclear binary mask was used to segment cell by the voronoi algorithm. In order to define ROIs the particle analyzer, a build-in FIJI plugin, was applied to the binary image of segmented cells. Although acquired images contain information of the subcellular activity we simplified image analysis pipeline in order be able to analyze 13900 images (more than 5000 cells) from a single experiment in a reasonable time window by averaging the intensity of FRET, CFP and FRET ratio over each ROI. However, significant compartmentalized signaling fluctuations might be averaged out. In all experiments, single-cell traces were normalized to the of the FRET ratio from before stimulation. The output file in a text format that was produced by FluoQ contained all measured parameters, statistical summaries.

### Outliers removal

In each independent experiment, single-cell traces were cautiously examined for artificial intensity spikes in the CFP and FRET channels. Those spikes were due to lamp intensity fluctuation, cell division, cell movement and, in rare cases, cell death. Therefore, we have developed the automated algorithm that detected artificial intensity spikes and removed cell traces containing those spikes from further data analysis. First, the baseline and response values were smoothed by running median (width of median window = 5). Subsequently, the difference between real values and smoothed values was used to calculated sample quantiles. The quantile mean and standard deviation for both baseline and response was calculated from the values belonging to the interval lying between 5^th^ and 95^th^ quantiles. Afterward, the z-score was calculated for each value of the baseline and response using quantile mean and standard deviation. Single cell traces containing z-score in the experimentally defined range (z-score ≥ 40 or z-score ≤ −40) were automatically removed. After removal single cell traces containing spikes, the mean baseline (formula 1), the standard deviation of the baseline (formula 2), the slope of the baseline (formula 3), the mean (formula 4), maximum (formula 5) and minimum (formula 6) response were calculated for each single-cell trace. Those features were subjected to the
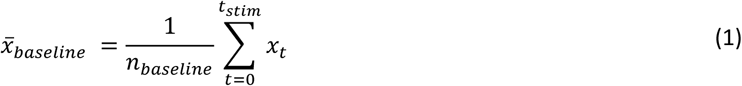

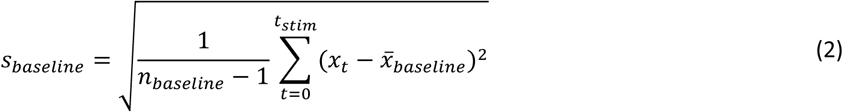

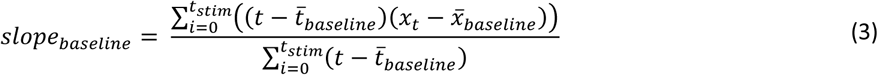

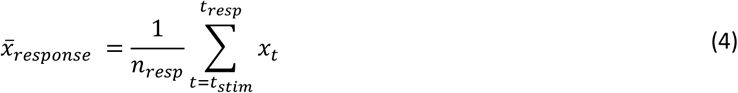

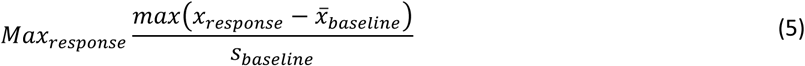

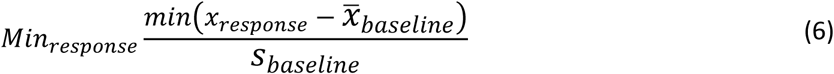

algorithm that classifies an outlier if it falls outside the interval defined in the following formula:

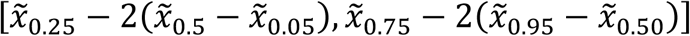

Once any feature of a particular single-cell trace was detected as an outlier, the cell was excluded from the analysis.

### Interaction Score

In order to calculate an interaction mode between two stimuli for each signaling node (or FRET biosensor), single-cell traces first were normalized by dividing each time point with the mean baseline 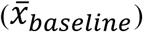 and with the mean value of untreated cells (x_mock) at each time point followed by a subtraction of 1. The resulted single-cell trace was normalized to the maximum observed average value across all conditions (formula 7). Afterwards, area under the curve (AUC) was computed for each single-cell trace within a specified time window (formula 8).
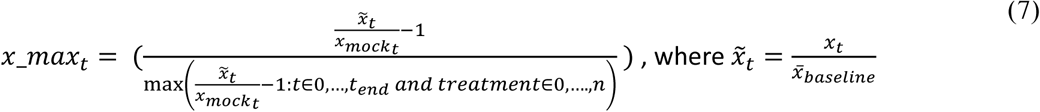

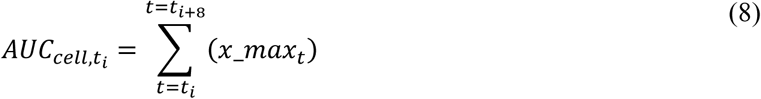

The expected response upon co-stimulation was calculated by the simple addition of the AUC mean of individual treatments. The combined error was estimated by an error propagation of the individual standard errors of the mean applying a Variance formula (Ku, 1966). In order compare the experimental and expected (calculated) additivity the means of two groups were subjected to Student’s two sample t-test and subsequently obtained P-values were corrected for multiple testing (Benjamini and Hochberg, 1995). We defined the ‘interaction score’ as the difference between the experimental and expected (calculated) additivity. In order to simplify visualization, the interaction score was scaled to the maximum interaction score observed for a FRET biosensor, giving a value that ranges from −1 (antagonism) to +1 (synergy). Adjusted p value < 0.05 was considered to be statistically significant.

### Analysis of RNA-seq data

RNA-seq reads were aligned with salmon version 0.8.2 (Patro, 2017) to the Human Reference Genome GRCh38 obtained from Ensembl (http://www.ensembl.org). Differential expression analysis was performed using the R/Bioconductor package DESeq2 (Love et al., 2014). To account for systematic effects between biological replicates, a blocking factor was added to the design formula prior to differential testing. Interaction analysis was carried out by supplying DESeq2 with a custom design matrix. The columns of the matrix correspond to factor levels of the respective EGF and IGF concentration and replicate as well as four columns corresponding to the interaction of the combined EGF/IGF treatment for the concentrations EGF:100ng/ml+IGF:100ng/ml, EGF:100ng/ml+IGF:6.25ng/ml, EGF:12.5ng/ml+IGF:12.5ng/ml and EGF:6.25ng/ml+IGF:100ng/ml respectively. Interaction scores were computed as model coefficient corresponding to the respective interaction column in the design matrix. Enrichment analysis was performed for genes with significant interactions using Fisher’s exact test with the R package piano (Varemo et al., 2013) on the Hallmark (Liberzon et al., 2015) gene set collection of MSigDB (Subramanian et al., 2005). Throughout the analysis, the false discovery rate (FDR) for calling differential expression or interaction was controlled using the method of Benjamini and Hochberg (Benjamini and Hochberg, 1995). If not otherwise stated, genes with an FDR < 0.1 were considered significant.

### Transcription factor binding sites analysis

Lists of 119 synergistic and 88 antagonistic genes (|GIS| >0.5 and adjusted p value <0.05) were uploaded to Pscan website (http://159.149.160.88/pscan/). The JASPAR non-redundant database (Jaspar 2018_NR) (Khan et al., 2018) and the promoter region between −450 and +50 bp of gene were used for motif analysis.

### General statistical analyses and FRET data visualization

Single-cell traces first were normalized by dividing each time point with respect to the mean baseline 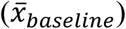 and with respect to the mean value of untreated cells (x_mock) at each time point followed by a subtraction of 1. The resulted single-cell traces were normalized to the maximum observed average value across all conditions (formula 7). The treated starved cells were normalized to starved “Mock” cells whereas treated cells kept in 10% FBS were normalized to untreated “Mock” cells maintained in 10% FBS. Unless otherwise stated, a FRET biosensor response is represented as normalized FRET ratio mean of all individual cells from identical conditions +/− SEM where the SEM is calculated from all cells in all experiments with identical conditions.

### Computation of correlation networks

To compute correlation networks, we first combined interaction score values across all time points, concentrations and ratios in a single vector for each FRET biosensor. For each growth factor pair, we used the Pearson correlation coefficient to quantify correlation between each pair of FRET biosensors. Correlation analyses were performed in R with Benjamini-Hochberg multiple testing correction separately within each condition. The correlation matrices were calculated by using the ‘rcorr’ function of Hmisc package (Harrell, 2016) and visualized by using R corrplot package. Network diagrams were generated using the igraph package (Csardi and Nepusz, 2006).

### Principal component analysis (PCA) and t-SNE analysis

The prcomp function without centering and scaling of the program R was used to perform PCA. The Rtsne package was used to perform t-SNE analysis. PCA and t-SNE was performed on X x Y matrix, where Y is different FRET biosensors each with X that included: time points, stimuli and stimuli doses.

### Data visualization

If not stated otherwise, the data was visualized using the R package ‘ggplot2’ (Wickham, 2009).

